# Temporal Dynamics of SMARCA4 Loss Define Its Role in Myogenic Differentiation of Alveolar Rhabdomyosarcoma

**DOI:** 10.64898/2026.10.20.760536

**Authors:** Charity Conlin Jennings, Emma Lashyro, JiaoJiao Wang, Gillian St John, Narendra Bharathy, Charles Keller

## Abstract

SMARCA4 (BRG1) is a critical ATPase subunit of canonical BAF complexes and a known long-term dependency in alveolar rhabdomyosarcoma (ARMS). To define the short-term consequences of SMARCA4 loss, we used a doxycycline-inducible shRNA system in Rh4 cells. Doxycycline treatment produced strong SMARCA4 knockdown, yet acute knockdown did not alter proliferation or induce myogenic differentiation. Ki-67 positivity remained unchanged, and Myosin Heavy Chain (MHC) expression was absent, indicating preservation of the undifferentiated, proliferative ARMS phenotype. These findings contrast with CRISPR-mediated *SMARCA4* knockout studies, which report cell-cycle exit and robust differentiation, suggesting that ARMS cells tolerate partial or early-phase SMARCA4 loss without destabilizing BAF complex function or PAX3-FOXO1–driven regulatory circuitry. Together with prior long-term knockdown studies, our results support a model in which SMARCA4 dependency in ARMS is a sustained phenomenon requiring deep, prolonged suppression to trigger differentiation and loss of tumor viability.

## INTRODUCTION

Rhabdomyosarcoma (RMS) is the most common pediatric soft tissue sarcoma, accounting for 5–10% of childhood malignancies [1-3]. Alveolar rhabdomyosarcoma (ARMS), defined by PAX3::FOXO1 or PAX7::FOXO1 fusion oncoproteins, carries the poorest prognosis, with five-year survival rates for metastatic disease below 10% despite multimodal therapy [4-8]. PAX3::FOXO1 establishes a core regulatory circuitry (CRC) that enforces proliferation and blocks terminal myogenic differentiation [9]. This blockade is sustained through cooperation with chromatin-remodeling machinery, particularly the SWI/SNF (BAF) family of ATP-dependent complexes, which regulate enhancer accessibility and lineage specification [10-13].

Our laboratory previously demonstrated that SMARCA4 (BRG1) is prominently expressed in ARMS, preferentially incorporated into canonical BAF complexes, and required for sustaining PAX3::FOXO1 expression and chemotherapy resistance [14]. CRISPR screening identified SMARCA4 as a long-term dependency: short-term knockdown produced modest effects, whereas sustained loss impaired growth, reduced tumor progression *in vivo*, and destabilized transcriptional programs maintaining the undifferentiated state [14, 15]. Complementary studies confirmed that SMARCA4-containing BAF complexes actively suppress differentiation in fusion-positive RMS, with complete SMARCA4 loss destabilizing CRC activity and inducing partial myogenic differentiation [16].

In this study, we evaluated the short-term consequences of *SMARCA4* knockdown in ARMS using a doxycycline-inducible shRNA system. Despite substantial reduction of SMARCA4 protein, Rh4 cells retained proliferative activity and showed no evidence of myogenic differentiation, as indicated by unchanged Ki-67 positivity and absent Myosin Heavy Chain (MHC) expression. These findings demonstrate that acute *SMARCA4* knockdown is insufficient to disrupt proliferation or relieve the differentiation blockade characteristic of ARMS. Our results align with prior work showing that differentiation phenotypes emerge only after sustained *SMARCA4* knockdown [16], suggesting that ARMS cells tolerate partial SMARCA4 loss without destabilizing canonical BAF complex function.

Importantly, pharmacologic inhibition of both SMARCA4 and SMARCA2 ATPase activity has been shown to produce a more rapid and pronounced differentiation response, with myogenic features emerging within 72 hours [16]. This highlights that simultaneous suppression of both ATPases—rather than SMARCA4 loss alone—may be required to fully overcome the differentiation blockade. Our findings therefore provide an initial framework for dissecting the temporal requirements of SMARCA4 function in ARMS and establish inducible knockdown systems as a platform for testing whether prolonged or dual ATPase targeting can drive meaningful myogenic reprogramming.

Together, these observations advance our understanding of SMARCA4 dependency in ARMS. These reinforce the concept that short-term, partial SMARCA4 loss preserves CRC activity and proliferation, while long-term, near-complete knockdown destabilizes enhancer landscapes, activates myogenic programs, and ultimately impairs tumor viability. This work informs the development of SMARCA4-targeted therapeutic strategies and underscores the importance of timing and completeness of ATPase suppression in overcoming the differentiation blockade in fusion-positive ARMS.

## METHODS

### Cell Culture and Doxycycline-Inducible Knockdown of *SMARCA4*

Human Rh4 rhabdomyosarcoma cells were cultured in RPMI-1640 medium supplemented with 10% fetal bovine serum (FBS; Cat# A5256501, Gibco, Thermo Fisher Scientific, Waltham, MA) and 1% penicillin–streptomycin (P/S; Cat# P4333, Sigma-Aldrich, St. Louis, MO) at 37 °C and 5% CO_2_. Cells were stably transduced with a lentiviral-mediated, doxycycline-inducible shRNA system to achieve targeted knockdown of *SMARCA4*. Lentiviral particles included three unique target-specific *SMARCA4* shRNAs and one scramble shRNA, all driven by a tet-inducible promoter construct (Cat# HSH-817261-100GFP, FencisBio, San Diego, CA).

The target sequences for the SMARCA4 shRNA constructs were:

- *SMARCA4* shRNA-1 (tet; Cat# HSH-817261-1-GFP, FencisBio): 5’-CCCGTGGACTTCAAGAAGATAA-3’
- *SMARCA4* shRNA-2 (tet; Cat# HSH-817261-2-GFP, FencisBio): 5’-CAAGCAAGATGTCGATGATGAA-3’
- *SMARCA4* shRNA-3 (tet; Cat# HSH-817261-3-GFP, FencisBio): 5’-TAGCAGCAGATGTAGTTTCAGA-3’

Among the three shRNA constructs tested, *SMARCA4*-targeting construct #1 (sh1) produced the most efficient knockdown and was therefore selected for all subsequent functional assays, including analyses of myogenic differentiation and proliferation. For knockdown experiments, cells were treated with doxycycline hydrochloride (Cat# D3072-1ML, Sigma-Aldrich) at 2 µg/mL for 5 days to induce shRNA expression, with untreated cells maintained in parallel as controls.

Human Skeletal Muscle Myoblasts (HSMM) were cultured as proliferating myoblasts in SkBM™-2 Basal Medium (Cat# CC-3246, Lonza, Walkersville, MD) supplemented with SkGM™-2 SingleQuots™ (Cat# CC-3244, Lonza) and the complete growth kit required for HSMM expansion (Cat# CC-3245, Lonza). HSMM were maintained under subconfluent conditions to preserve the myoblast state. For differentiation assays, HSMM were switched at ~70% confluence to DMEM/F-12 supplemented with 2% horse serum (Cat# 31874, Invitrogen, Thermo Fisher Scientific) and maintained until robust myogenic differentiation was observed.

### Simple Western (Jess) Protein Quantification

SMARCA4 protein levels were measured using the Jess Simple Western system (ProteinSimple, Bio-Techne, San Jose, CA) according to the manufacturer’s instructions. Rh4 cells were harvested after 5 days of doxycycline treatment, and total protein lysates were prepared using RIPA Lysis and Extraction Buffer (Cat# 89900, Thermo Scientific). Protein concentrations were determined using the Pierce™ BCA Protein Assay Kit (Cat# 23225, Thermo Scientific), and equal amounts of protein were loaded into 12–230 kDa Separation Module plates (Cat# SM-W001, ProteinSimple) for automated capillary-based immunodetection.

Primary antibodies included anti-SMARCA4/BRG1 (1:100; clone G-7, Cat# sc-17796, Santa Cruz Biotechnology, Dallas, TX) and anti-β-actin (1:25; clone mAbcam 8226, Cat# ab8226, Abcam, Cambridge, MA). HRP-conjugated secondary antibodies (Cat# 042-205 and 042-206, ProteinSimple) and chemiluminescent reagents were processed automatically by the Jess instrument. Digital signals were quantified using Compass for Simple Western software (ProteinSimple). SMARCA4 peak areas were normalized to β-actin and expressed relative to untreated controls.

Electrophoretic separation, immunoprobing, and signal detection were carried out within the automated capillary system. Digital chemiluminescent signals were quantified using Compass for Simple Western software (ProteinSimple). SMARCA4 peak areas were normalized to β-actin and expressed relative to untreated controls.

### Immunocytochemistry and Quantification of Ki-67 Positive Cells

Rh4 cells expressing control or *SMARCA4*-targeting shRNA were treated with doxycycline for 5 days prior to analysis. Cells were fixed and stained for Ki-67 using the Ki-67 Monoclonal Antibody (SolA15), Alexa Fluor™ 488 (Cat# 53-5698-82, eBioscience™, Invitrogen, Thermo Fisher Scientific). HSMM differentiated in DMEM/F-12 with 2% horse serum served as a negative control population.

Following fixation and permeabilization, samples were incubated with the primary antibody at 1:200, washed, and counterstained with Hoechst nuclear dye (Invitrogen, Thermo Fisher Scientific). Images were acquired using a Zeiss LSM confocal microscope (Carl Zeiss Microscopy, LLC, White Plains, NY).

Ki-67 positivity was calculated as the number of Ki-67^+^ nuclei divided by total nuclei per field. Multiple fields were analyzed for each biological replicate. The percentage of Ki-67^+^ cells in shControl samples was normalized to 100%. Data are presented as mean ± SD (n = 3), and statistical significance was evaluated using an unpaired two-tailed Student’s t-test in GraphPad Prism 10 (Order Code PRISM-10, GraphPad Software, LLC, San Diego, CA).

### Immunocytochemistry for MHC

For detection of terminal myogenic differentiation, cells were fixed after 5 days of doxycycline treatment and stained for Myosin Heavy Chain (MHC) using the MF-20 monoclonal antibody (Developmental Studies Hybridoma Bank, DSHB, Iowa City, IA) at 1:100. After primary antibody incubation and washes, samples were incubated with Goat anti-Mouse IgG (H+L) Cross-Adsorbed Secondary Antibody, Cyanine-5 (Cat# A10524, Invitrogen) at the manufacturer-recommended dilution.

Nuclei were counterstained with Hoechst 33258 (Cat# H3569, Invitrogen, Thermo Fisher Scientific) at 1:1000, and images were acquired using a Zeiss LSM confocal microscope (Carl Zeiss Microscopy). HSMM differentiated in DMEM/F-12 with 2% horse serum served as positive controls.

Quantification of MHC-positive cells was performed by calculating the ratio of nuclei within MHC^+^ myotubes to total nuclei. The percentage of MHC^+^ cells in control samples was normalized to 100%. For each condition, at least 600 nuclei were counted across three biological replicates. Data are presented as mean ± SD (n = 3), and statistical significance was assessed using Student t test in GraphPad Prism 10 (GraphPad Software).

### Statistical Analysis

In most experiments, statistical significance was assessed using Student’s t test, with *P* < 0.05 considered significant. Significance thresholds were defined as *P* < 0.05 (\**) P* < 0.01 (**) and *P* < 0.001 (***). Error bars represent the mean ± standard deviation (SD).

## RESULTS

### Doxycycline efficiently induces *SMARCA4* knockdown in Rh4 cells

To evaluate the consequences of *SMARCA4* loss in alveolar rhabdomyosarcoma (ARMS), we first confirmed the effectiveness of our doxycycline-inducible shRNA system in Rh4 cells. Immunoblot analysis demonstrated a marked reduction in SMARCA4 protein abundance after five days of doxycycline exposure at 2 µg/mL (Fig. 1A). Densitometric quantification across biological replicates verified a consistent and significant decrease in SMARCA4 expression relative to untreated controls (Fig. 1B). These data established that doxycycline induction reliably reduces SMARCA4 in Rh4 cells.

**Figure 1.**
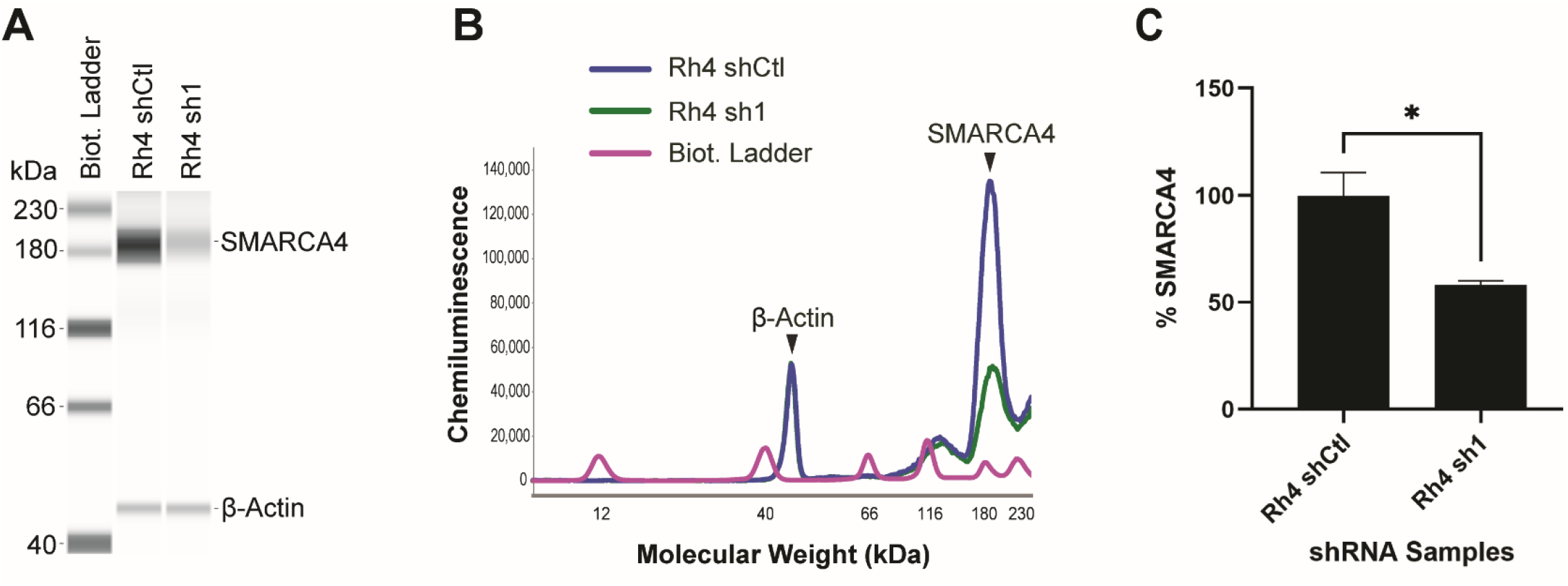
Doxycycline-inducible knockdown of *SMARCA4* in Rh4 cells. (A) Immunoblot analysis confirming reduction of SMARCA4 protein levels after 5 days of doxycycline treatment in Rh4 cells expressing *SMARCA4*-targeting shRNA (sh1) compared with scrambled control (shCtl). β-actin was used as a loading control. Representative blot from *n* = 3 independent experiments. (B) Chemiluminescence electropherogram traces illustrating decreased SMARCA4 signal intensity in sh1 relative to shCtl, with overlapping β-actin peaks unchanged. (C) Densitometric quantification of SMARCA4 protein levels normalized to β-actin and expressed relative to untreated controls. Quantification confirms a reproducible reduction in SMARCA4 abundance following doxycycline treatment. Data represent mean ± SD from *n* = 3 biological replicates. Statistical significance was assessed using Student’s t test in GraphPad Prism, *P* < 0.05 (*).

### SMARCA4 loss does not impair short-term proliferation in Rh4 cells

We next examined whether SMARCA4 knockdown affects proliferative capacity. Immunocytochemical detection of Ki-67 revealed no observable difference in Ki-67 expression between control and *SMARCA4*-knocked-down Rh4 cells following five days of doxycycline treatment (Fig. 2A). The proportion of Ki-67–positive nuclei remained comparable between groups, indicating that *SMARCA4* knockdown does not diminish short-term proliferation. As expected, differentiated human skeletal muscle myoblasts (HSMM) exhibited low Ki-67 positivity and served as a negative control. Quantification of Ki-67–positive cells corroborated these observations, showing no reduction in proliferative fraction upon *SMARCA4* knockdown (Fig. 2B). Together, these findings demonstrate that SMARCA4 knockdown does not impair Rh4 cell proliferation under short-term conditions.

**Figure 2.**
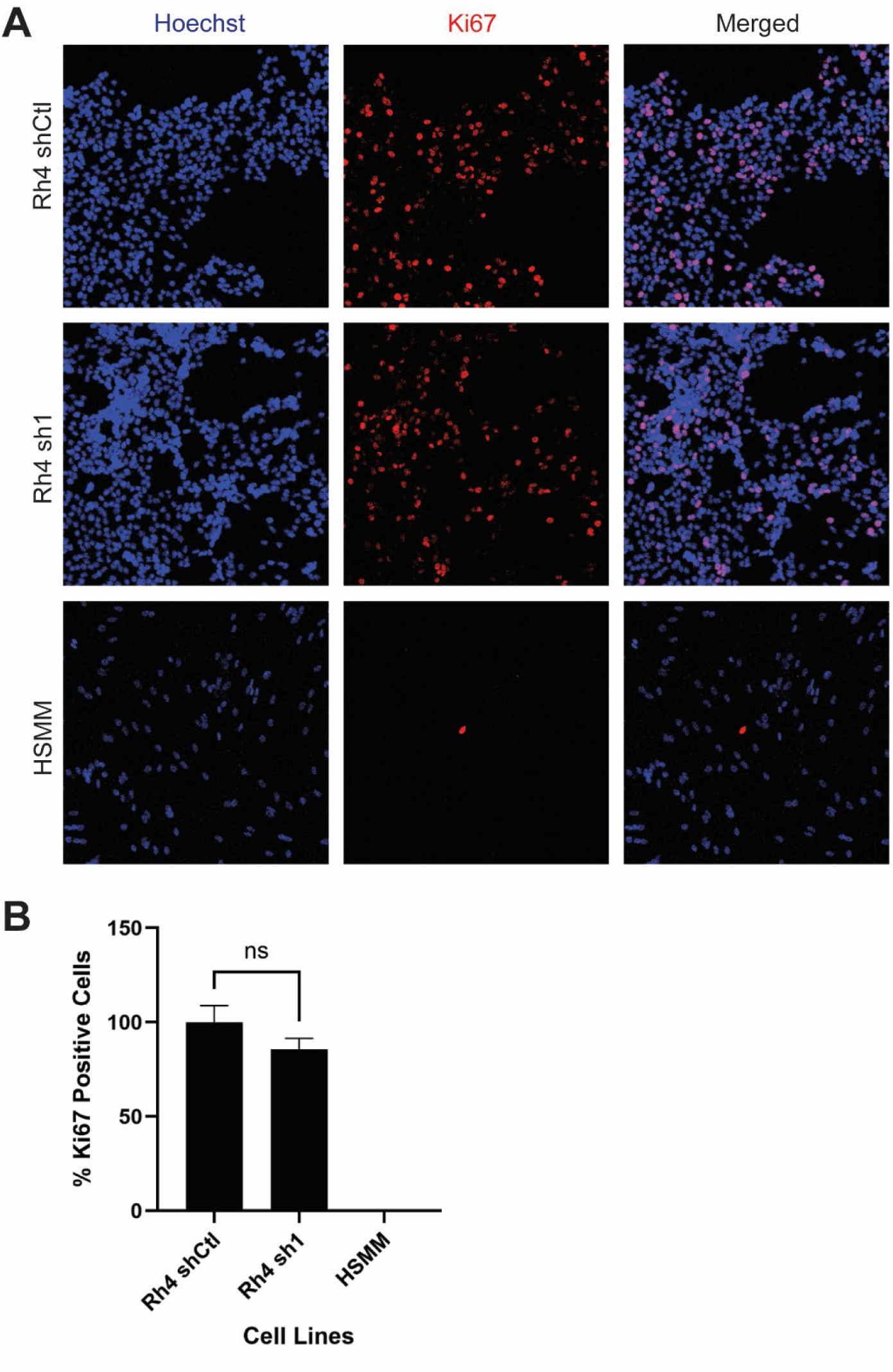
Knockdown of *SMARCA4* does not reduce proliferation in Rh4 cells. (A) Immunocytochemical detection of Ki-67 in control and *SMARCA4*-knocked-down Rh4 cells following 5 days of doxycycline treatment. SMARCA4 loss does not alter Ki-67 expression, and the proportion of Ki-67–positive nuclei remain comparable between groups. Human Skeletal Muscle Myoblasts (HSMM) cells were included as negative control as myoblasts exit cycle and stop proliferating upon serum starvation-induced differentiation. Representative images from *n* = 3 independent experiments. (B) Quantification of Ki-67–positive cells (percentage of Ki67-positive cells/total cells) in control and *SMARCA4*-knocked-down Rh4 cells. The percentage of Ki-67^+^ cells in ShCtrl were given a value of 100%. Consistent with the imaging results, *SMARCA4* knockdown does not reduce proliferation. Data represent mean ± SD from *n* = 3 biological replicates. Statistical significance was assessed using an unpaired two-tailed Student’s t-test in GraphPad Prism. ns, *P* > 0.05, where ns denotes not significant.

### SMARCA4 knockdown does not induce myogenic differentiation

Given the role of SMARCA4 in chromatin remodeling and lineage specification, we assessed whether SMARCA4 loss promotes myogenic differentiation. Immunofluorescence staining for Myosin Heavy Chain (MHC) revealed no detectable MHC expression in *SMARCA4*-knocked-down Rh4 cells (Fig. 3A). Morphologically, SMARCA4-depleted cells did not exhibit features characteristic of terminal myogenic differentiation (elongation, multinucleation, withdrawal from cell cycle division). In contrast, HSMM cells displayed robust MHC expression and served as positive control. Quantification confirmed the absence of MHC-positive cells in both control and SMARCA4-knocked-down Rh4 populations (Fig. 3B). These results indicate that SMARCA4 knockdown alone is insufficient to initiate terminal myogenic differentiation in Rh4 cells.

**Figure 3.**
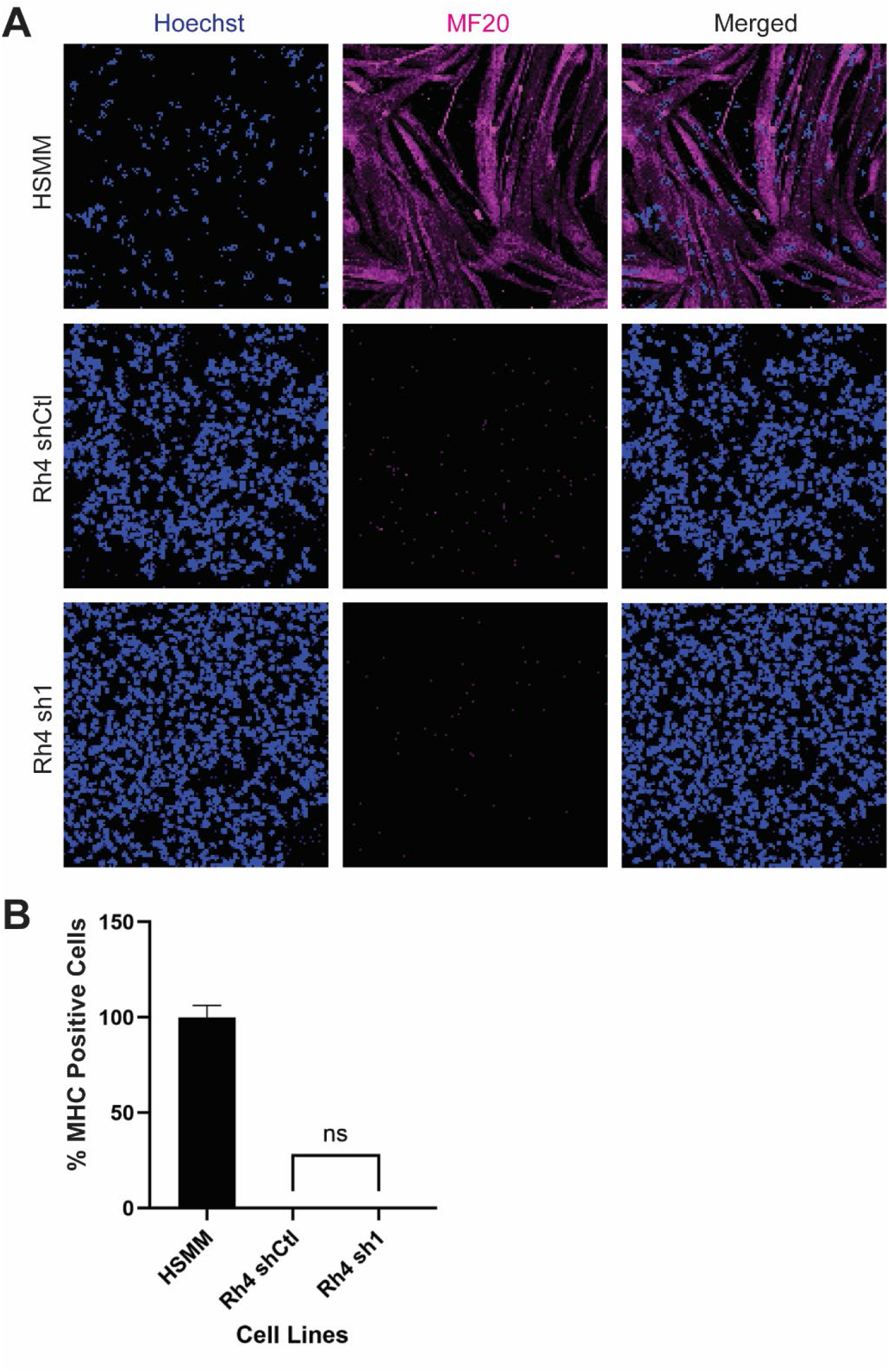
Knockdown of *SMARCA4* does not induce meaningful myogenic differentiation over 5 days. (A) Immunofluorescence staining for myogenic differentiation markers in control and SMARCA4-knocked-down cells. Loss of SMARCA4 does not result in detectable expression of the terminal myogenic marker Myosin Heavy Chain (MHC), and *SMARCA4*-knocked-down cells do not exhibit morphological features of myogenic differentiation. HSMM were included as positive control. Representative images from *n* = 3 independent experiments. (B) Quantification of MHC-positive cells in control versus *SMARCA4*-knocked-down cells after 5 days of doxycycline treatment. Consistent with the imaging results, *SMARCA4* loss does not induce myogenic differentiation, as reflected by no MHC-positive cells. The percentage of MHC-positive cells in HSMM was set to 100%. Data represent mean ± SD from *n* = 3 biological replicates. Data represent mean ± SD from *n* = 3 biological replicates. Statistical significance was assessed using Student’s t test in GraphPad Prism. ns, *P* > 0.05, where ns denotes not significant.

## DISCUSSION

In this study, we evaluated the short-term consequences of *SMARCA4* (*BRG1*) knockdown in alveolar rhabdomyosarcoma (ARMS) using a doxycycline-inducible shRNA system. Although SMARCA4 protein levels were substantially reduced, Rh4 cells retained normal proliferative activity and exhibited no evidence of myogenic differentiation. Ki-67 positivity remained unchanged, and Myosin Heavy Chain (MHC) expression was undetectable, indicating that acute SMARCA4 loss is insufficient to disrupt the proliferative state or relieve the differentiation blockade characteristic of ARMS. These findings provide an initial framework for understanding SMARCA4 dependency in Rh4 cells and establish the foundation for extending inducible knockdown analyses across additional ARMS models.

Our observations add context to the differentiation phenotype reported by Laubscher *et al* [16]. In that study, CRISPR-mediated SMARCA4 knockout in fusion-positive rhabdomyosarcoma over seven days induced modest cell cycle exit and partial myogenic differentiation. Several mechanistic considerations and differences likely explain this discrepancy. First, CRISPR knockout produces abrupt and near-complete elimination of SMARCA4, whereas shRNA-mediated knockdown results in a gradual and sometimes incomplete reduction. ARMS cells appear capable of tolerating partial SMARCA4 loss without destabilizing canonical BAF complex function [17]. Second, the differentiation phenotype described by Laubscher *et al* emerges only after sustained *SMARCA4* knockdown; our five-day induction window may capture only the early phase of knockdown, prior to epigenetic destabilization. Third, complete *SMARCA4* loss disrupts the CRC driven by PAX3-FOXO1, MYCN, MYOD1, and MYOG, enabling MYCN enhancer invasion and activation of myogenic super-enhancers. In contrast, residual SMARCA4 in our system may maintain CRC stability. Finally, *SMARCA4* knockout permits compensatory redistribution of BRM-containing complexes to myogenic loci, a process that may not occur under partial knockdown.

A closer examination of the Laubscher study further supports this interpretation. Although SMARCA4 loss produced detectable myogenic marker induction, the extent of differentiation appeared modest, with limited evidence of robust MHC upregulation and only partial activation of late myogenic programs. Cell cycle effects were similarly restrained, consistent with only a mild reduction in proliferative activity after prolonged SMARCA4 knockdown. Notably, pharmacologic inhibition or degradation of both SMARCA4 and SMARCA2 ATPase activity produced a more rapid and pronounced differentiation response, with myogenic features emerging within 72 hours [16]. This observation suggests that simultaneous suppression of both ATPases—rather than SMARCA4 loss alone may be required to fully overcome the differentiation blockade in fusion-positive ARMS. Together, these observations indicate that complete and sustained disruption of SWI/SNF ATPase function is necessary to drive meaningful myogenic reprogramming in this disease.

Importantly, our results are fully consistent with our prior publication, which demonstrated that short-term *SMARCA4* knockdown produces modest phenotypic effects, whereas long-term SMARCA4 loss is incompatible with ARMS survival [14], not myogenic differentiation. In our prior study, CRISPR screens, colony formation assays, and *in vivo* models revealed a strong dependency on SMARCA4 for sustained tumor growth. Bharathy *et al* [14], further showed that SMARCA4 is preferentially incorporated into canonical BAF complexes and supports PAX3-FOXO1–driven transcriptional programs, reinforcing the concept that ARMS cells rely on SMARCA4 to maintain epigenetic stability [14]. Our current findings align with this model: short-term knockdown does not impair proliferation or induce differentiation, underscoring that SMARCA4 dependency in ARMS is a long-term phenomenon requiring deep and sustained knockdown to manifest.

Collectively, these observations support a unified mechanistic interpretation. Short-term, partial SMARCA4 loss preserves BAF complex integrity and CRC activity, allowing ARMS cells to maintain proliferation and an undifferentiated state. In contrast, long-term, near-complete SMARCA4 loss destabilizes BAF complexes, rewires enhancer landscapes, activates MYOD1 and MYOG, and ultimately triggers terminal myogenic differentiation and loss of tumor viability. The inducible shRNA systems developed here provide the experimental framework necessary to determine whether prolonged or more complete *SMARCA4* suppression can reproduce the differentiation phenotypes observed with CRISPR knockout and whether dual ATPase targeting (SMARCA4/SMARCA2) is required to overcome the differentiation blockade in fusion-positive ARMS [16].

A notable limitation of this work is that all short-term knockdown experiments were conducted exclusively in Rh4 cells. This choice was intentional: Rh4 is the ARMS model in which Laubscher *et al* most clearly demonstrated SMARCA4 loss-induced myogenic differentiation, making Rh4 the most appropriate line for initial comparison. However, reliance on a single cell line limits generalizability. ARMS models differ in transcriptional wiring, MYCN activity, and BAF complex composition, and other lines may exhibit distinct responses to *SMARCA4* knockdown.

Ongoing studies using inducible knockdown systems across Rh5, Rh30, and Rh41 will address this limitation and determine whether the short-term tolerance to SMARCA4 loss observed in Rh4 is universal or cell line-specific. Parenthetically, a SMARCA4 PROTAC that leaves SMARCA2 intact would be a tool of great interest for dissecting the early, intermediate, and late time scale of near-complete SMARCA4 knockdown.

In summary, this work advances our understanding of SMARCA4 dependency in ARMS and informs the development of SMARCA4-targeted therapeutic strategies. Determining whether genetic or pharmacologic suppression of SMARCA4 can induce differentiation and suppress tumor growth will be essential for translating these mechanistic insights into new treatment approaches for this aggressive pediatric malignancy.

## ACKNOWLEDGEMENTS

Funding for this work was supported by NIH R01CA258720 and donors honoring Maya Handler.

## AUTHORS’ CONTRIBUTIONS

Overall conception and design: CK

Analysis and interpretation of experimental data: CK, NB, CJ, JJ, EL

Development of experimental methodology: CK, NB

Acquisition of experimental data: CJ, JJ, GSJ

Statistical analysis: CJ

Writing, review and/or revision: CK, NB

Study supervision: CK

## LIST OF ABREVIATIONS

RMS: rhabdomyosarcoma
aRMS: alveolar rhabdomyosarcoma
CRC: core regulatory circuitry
MHC: myosin heavy chain

